# The neutralization potency of anti-SARS-CoV-2 therapeutic human monoclonal antibodies is retained against novel viral variants

**DOI:** 10.1101/2021.04.01.438035

**Authors:** Efi Makdasi, Anat Zvi, Ron Alcalay, Tal Noy-Porat, Eldar Peretz, Adva Mechaly, Yinon Levy, Eyal Epstein, Theodor Chitlaru, Ariel Tennenhouse, Moshe Aftalion, David Gur, Nir Paran, Hadas Tamir, Oren Zimhony, Shay Weiss, Michal Mandelboim, Ella Mendelson, Neta Zuckerman, Ital Nemet, Limor Kliker, Shmuel Yitzhaki, Shmuel C. Shapira, Tomer Israely, Sarel J. Fleishman, Ohad Mazor, Ronit Rosenfeld

## Abstract

A wide range of SARS-CoV-2 neutralizing monoclonal antibodies (mAbs) were reported to date, most of which target the spike glycoprotein and in particular its receptor binding domain (RBD) and N-terminal domain (NTD) of the S1 subunit. The therapeutic implementation of these antibodies has been recently challenged by emerging SARS-CoV-2 variants that harbor extensively mutated spike versions. Consequently, the re-assessment of mAbs, previously reported to neutralize the original early-version of the virus, is of high priority.

Four previously selected mAbs targeting non-overlapping epitopes, were evaluated for their binding potency to RBD versions harboring individual mutations at spike positions 417, 439, 453, 477, 484 and 501. Mutations at these positions represent the prevailing worldwide distributed modifications of the RBD, previously reported to mediate escape from antibody neutralization. Additionally, the *in vitro* neutralization potencies of the four RBD-specific mAbs, as well as two NTD-specific mAbs, were evaluated against two frequent SARS-CoV-2 variants of concern (VOCs): (i) the B.1.1.7 variant, emerged in the UK and (ii) the B.1.351 variant, emerged in South Africa. Variant B.1.351 was previously suggested to escape many therapeutic mAbs, including those authorized for clinical use. The possible impact of RBD mutations on recognition by mAbs is addressed by comparative structural modelling. Finally, we demonstrate the therapeutic potential of three selected mAbs by treatment of K18-hACE2 transgenic mice two days post infection with each of the virus strains.

Our results clearly indicate that despite the accumulation of spike mutations, some neutralizing mAbs preserve their potency against SARS-CoV-2. In particular, the highly potent MD65 and BL6 mAbs are shown to retain their ability to bind the prevalent novel viral mutations and to effectively protect against B.1.1.7 and B.1.351 variants of high clinical concern.

## Introduction

An unprecedented worldwide research and development effort has resulted in the rapid development of several prophylactic and therapeutic immune tools to combat the COVID-19 pandemic caused by SARS-CoV-2. These tools predominantly target the virus spike glycoprotein which is essential for the attachment of the virus to the target cell and hence plays an essential role in virus infectivity (Walls et al., 2020). Emergency-authorized vaccines against the SARS-CoV-2 spike produced by Pfizer/BioNTech, Moderna, AstraZenica, Johnson & Johnson (Krammer, 2020) and others, are already being used in mass vaccination campaigns (https://www.who.int/publications/m/item/draft-landscape-of-covid-19-candidate-vaccines). Additionally, passive immunity was achieved by the administration of convalescent plasma or recombinant neutralizing monoclonal antibodies [mAbs; (Alam et al., 2021; Weinreich et al., 2021; Wu et al., 2020b)]. This therapeutic avenue accelerated the development of many potent neutralizing mAbs, primarily targeting the receptor binding domain (RBD) and the N-terminal domain (NTD) of the spike-S1 subunit [reviewed by (Xiaojie et al., 2020)]. A single therapeutic mAb, generated by Eli Lilly and Company, and a dual antibody combination, generated by Regeneron Pharmaceuticals, recently received emergency-use authorization (Chen et al., 2021a; Weinreich et al., 2021).

Prior to its global expansion, SARS-CoV-2 was expected to exhibit a relatively low mutations rates, as compared to many other RNA viruses since its genome encodes a proofreading exoribonuclease (Robson et al., 2020). Nevertheless, the long-term global spread of the SARS-CoV-2, possibly combined with selective pressure for immune escape (Kemp et al., 2021), enabled the emergence of new SARS-CoV-2 variants. Specifically, multiple mutations in the spike glycoprotein are evolving, including mutations that are located in the spike S1 subunit, particularly residing in the antigenic supersite of the NTD (Cerutti et al., 2021; McCallum et al., 2021; Noy-Porat et al., 2021) or in the RBD [hACE2-binding site; (Baum et al., 2020; Chen et al., 2020; Noy-Porat et al., 2020)], sites that represent a major target of potent virus-neutralizing antibodies.

The impact of accumulated mutations is closely monitored; yet, only a minor fraction, which are selectively favorable, might spread and reach high frequency, and more importantly, become fixed in the population. Emergence of such genetic variants has important epidemiological consequences since they may exhibit increased transmissibility, reinfection of vaccination or convalescent individuals, or increase disease severity. The WHO has recently established the working definitions of “SARS-CoV-2 Variant of Interest” (VOI) and of “SARS-CoV-2 Variant of Concern” (VOC) (https://www.who.int/publications/m/item/weekly-epidemiological-update---23-february-2021). One of the major VOCs identified and monitored recently, is denoted as 20I/501Y.V1 belonging to the B.1.1.7 lineage, which has a total of 18 nonsynonymous mutations relative to the original Wuhan strain. In this variant,7 replacements and 2 deletions reside in the spike protein [see Supplementary Figure 1 for schematic presentation; (Rambaut et al., 2020b)]. Since its first emergence in the UK in September 2020 (Rambaut et al., 2020a), the B.1.1.7 variant is rapidly globally spreading. As of June 2021, the variant has been detected in over 140 countries, with an apparent cumulative prevalence of 44% worldwide (for instance, 58%, 33% and 68% in the UK, US and Israel, respectively) and a worldwide average daily prevalence of ~75% (https://outbreak.info). Two additional VOCs were reported: the B.1.351 lineage (also known as 20H/501Y.V2; schematically depicted in Supplementary Figure 1), identified for the first time in October 2020 in South Africa (Tegally et al., 2021) and the P.1 lineage, (also known as 501Y.V3), first identified in December 2020 in Brazil (Faria et al., 2021). Both variants are less abundant worldwide (up to 2%) and mostly contained in the geographic surrounding of their originating site. The most recent variant determined by the WHO as VOC is the B.1.617.2 lineage, first identified in India, with an apparent cumulative prevalence of 3% worldwide (as of June 2021), and very recently also detected in Israel. This variant, harboring 9 mutations in the spike protein (among which L452R and T478K are in the RBD), is associated with higher transmissibility (Saito et al., 2021) and potential immune escape (Planas et al., 2021; Yadav et al., 2021). The full biological and clinical implications of the new SARS-CoV-2 variants are yet to be determined. Nevertheless, the careful immunological assessment of known mutations, in particular in the RBD, is essential, due to the possible impact on vaccines and therapeutic countermeasures, such as monoclonal antibodies. Of the multitude of possible genomic loci, mutations at several positions were already reported at relatively high frequency in the ~2×10^6^ sequences available to date (GISAID initiative, https://gisaid.org (Elbe and Buckland-Merrett, 2017). The most frequent mutation, N501Y, representing the hallmark of three circulating VOCs (B.1.1.7, B.1.351 and P.1), was first detected in February 2020, and as of June 2021 is present in over 70% of the global cases in more than 160 countries. The mutation S477N, was reported in 43% of the cases worldwide, since its emergence in February 2020. Its cumulative prevalence is most prominent in Australia (56% as of June 2021). The mutation E484K, has been detected in more than 120 countries, exhibiting a worldwide cumulative prevalence of 6%. This mutation was detected in the South African (B.1.351) and Brazilian (P.1) variants and recently in a UK “B.1.1.7+E484K” variant. The N439K, a sentinel receptor binding motif mutation (Thomson et al., 2021) has an apparent worldwide cumulative prevalence of 2%, reported in at least 79 countries. This mutation has emerged in multiple SARS-CoV-2 clades, and is mostly associated with the B.1.258 lineage derivatives, circulating in central Europe. The K417N mutation was reported in over 1% of the cases worldwide in at least 105 countries. This mutation represents one of the hallmarks of the B.1.351 lineage and is exhibited in approximately 50% of South African cases. The replacement Y453F was detected in at least 15 countries, predominantly in Denmark. In late 2020, this mutation raised substantial concern, when it was detected in a variant found in the mink population (Thomson et al., 2021).

Both predictive theoretical and experimental approaches revealed that escape mutants can rapidly occur when SARS-CoV-2 is exposed to selective pressures mediated by neutralizing polyclonal sera or individual mAbs (Andreano et al., 2020; Liu et al., 2021; Starr et al., 2021a; Weisblum et al., 2020). More specifically, escape mutations within the RBD were predicted and experimentally confirmed to affect its function (mainly with respect to hACE2 binding) and recognition by mAbs. Substitutions N501Y, E484K, K417N, Y453F, N439K and S477N, were among the most frequent mutations that mediated immune escape and were shown to reduce and even completely abrogate the neutralizing activity of several highly potent mAbs, including those which are already in clinical use (Andreano et al., 2020; Chen et al., 2021b; Liu et al., 2021; Starr et al., 2021a; Weisblum et al., 2020). These substitutions naturally occurred in infected individuals, most of which are now represented by SARS-CoV-2 emerged genetic variants which spread worldwide.

We previously reported the isolation of RBD- and NTD-specific mAbs (Barlev-Gross et al., 2021; Noy-Porat et al., 2020; Noy-Porat et al., 2021; Rosenfeld et al., 2021), among which the MD65 mAb showed exceptional neutralization potency, as demonstrated by *in vitro* and *in vivo* experiments.

In the current study, we present the re-evaluation of four SARS-CoV-2 neutralizing mAbs (MD65, MD62, MD29 and BL6), directed against four distinct epitopes in the spike RBD, for their ability to bind RBD variants that represent individual substitutions encountered in the VOCs. Additionally, we assessed the *in vitro* neutralization capacity of these four anti-RBD and two anti-NTD mAbs to counteract the SARS-CoV-2 B.1.1.7 and B.1.351 variants. Comparative structural modeling was conducted to determine the possible impact of mutations on the binding efficiency of the MD65 mAb. Finally, we evaluated the *in vivo* therapeutic potential of three selected mAbs by treatment of K18-hACE2 transgenic mice two days post infection with each of the virus strains.

## Results and Discussion

### Binding SARS-CoV-2 single mutated-RBD versions by specific mAbs

In the current study, we re-evaluated the antigen-binding of the recently reported mAbs, MD65, MD62, MD29 and BL6, targeting four distinct RBD epitopes [(Noy-Porat et al., 2020); Supplementary Figure 2]. The binding capability of these mAbs was tested with respect to six individual mutations in the SARS-CoV-2 spike recombinant RBD (rRBD), identified in circulating variants, including VOCs. These inspected mutations are N501Y, S477N, E484K, N439K, K417N and Y453F as detailed below (Figure 1A; for complete lineages and mutation reports, see https://outbreak.info).

**Figure 1.**
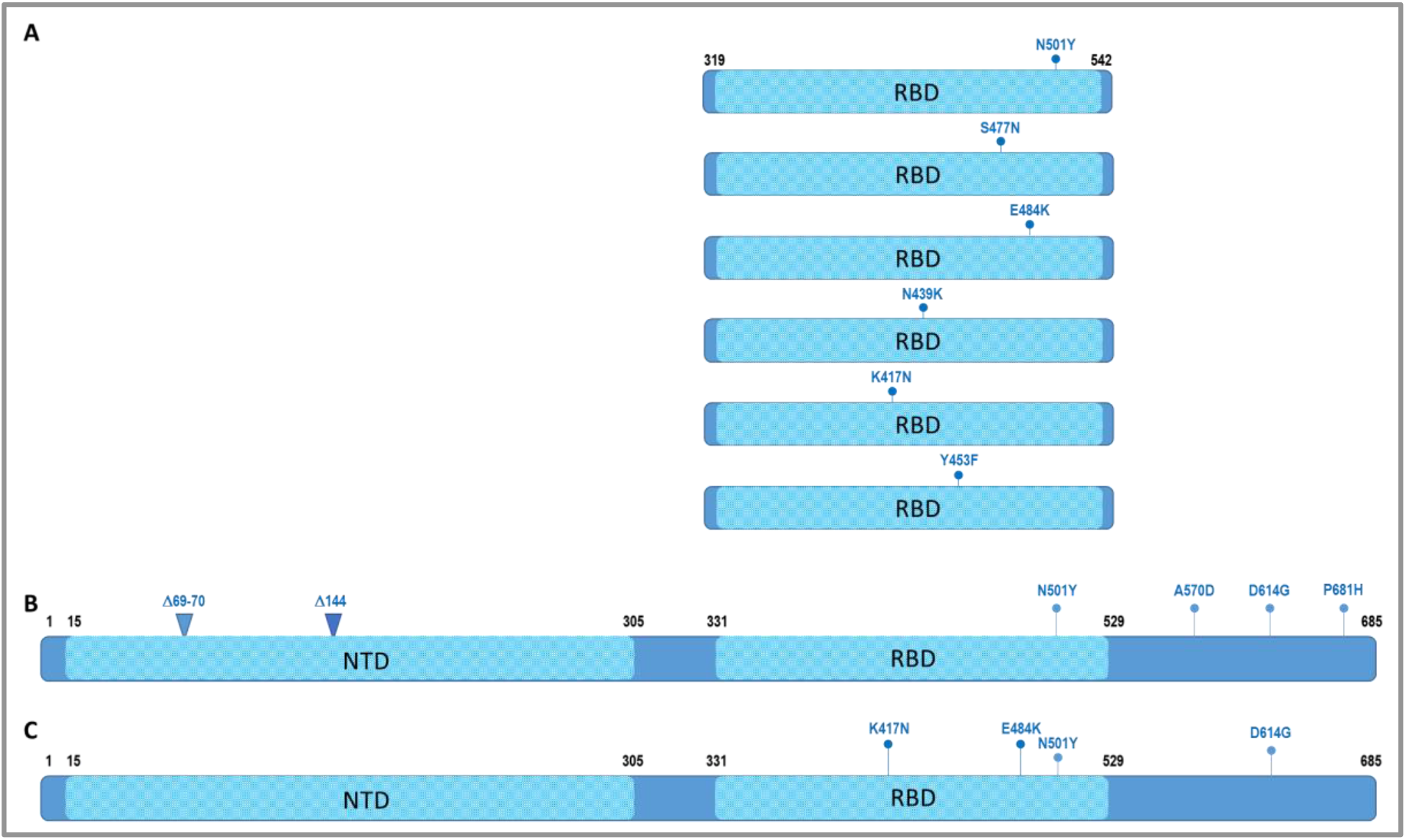
Schematic representation of SARS-CoV-2 RBD and S1 variants. **A**. Depiction of recombinant RBD variant proteins, each including the indicated single highly frequent replacements reported in the RBD domain. The domain coordinates are according to the recombinant RBD used throughout the study. **B**. Schematic representation of the spike S1 subunit, along with the replacements and deletions characterizing the SARS-CoV-2 B.1.1.7 genetic variant. **C**. Schematic representation of the spike S1 subunit, along with the RBD replacements, characterizing the SARS-CoV-2 B.1.351 genetic variant. The numbering is according to the Wuhan reference sequence (Accession no. NC_045512). For the full panel of replacements in the B.1.1.7 and B.1.351 spike protein, see Supplementary Figure 1.

Biolayer interferometry (BLI) analysis was applied to evaluate the ability of the four RBD-specific mAbs that we have previously reported to bind the SARS-CoV-2 single mutated-RBD variants. As presented in Figure 2, the binding of these mAbs was only slightly affected (5-22% loss of binding) by five of the six substitutions in the RBD. The only significant reduction in binding capacity, compared to the WT rRBD, was observed for the K417N mutant by the MD62 mAb (~74% reduction) and to a lesser extent by the MD65 mAb (17% reduced binding). In light of these results, it is anticipated that these mAbs, previously shown to neutralize SARS-CoV-2 by targeting distinct epitopes on the RBD, maintain their potency against variant strains carrying these mutations.

**Figure 2.**
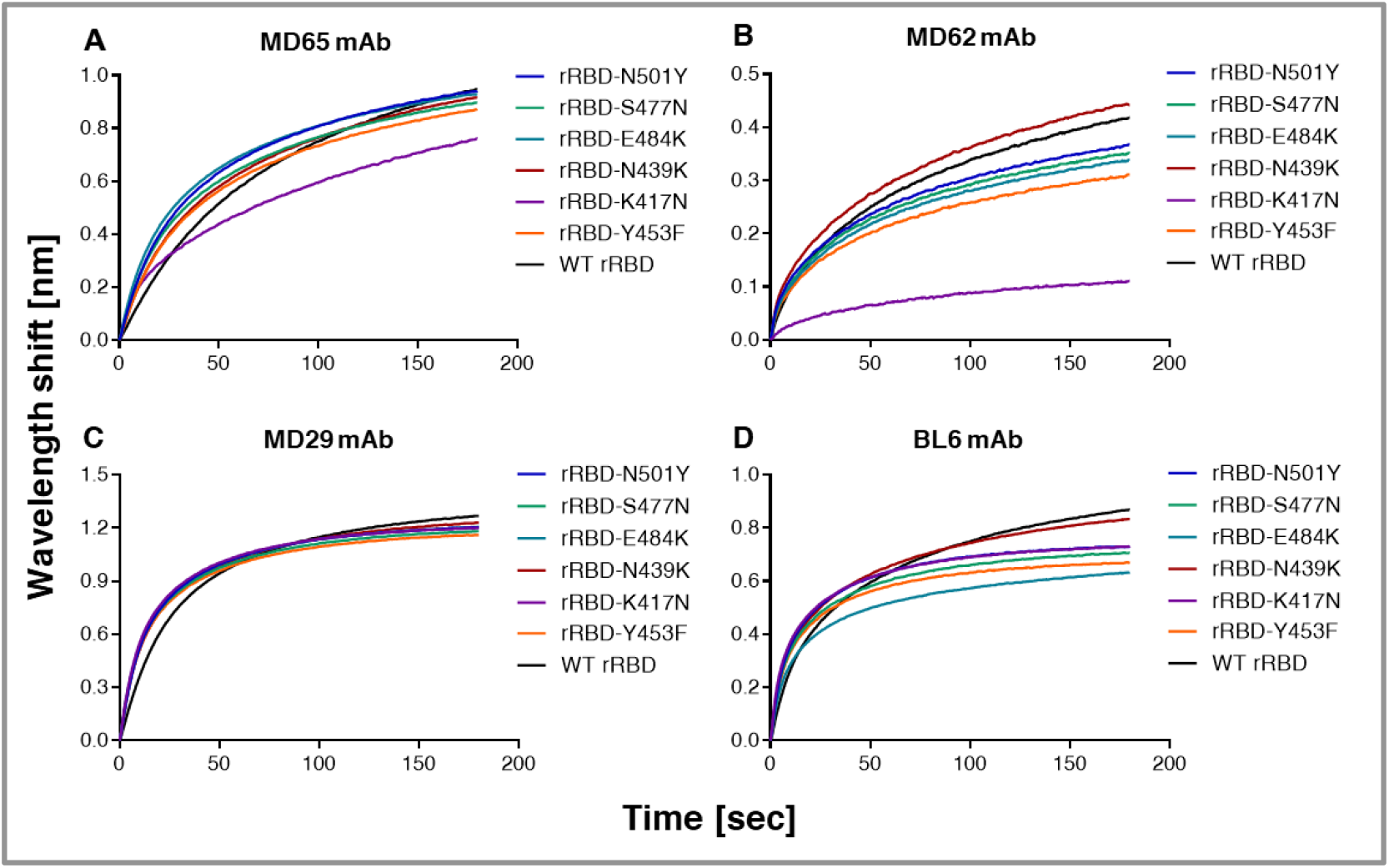
MAbs binding of singly-mutated rRBDs. Binding of the WT and the six indicated singly-mutated recombinant RBDs by MD65 (**A**), MD62 (**B**), MD29 (**C**) and BL6 (**D**) mAbs, was evaluated by Biolayer interferometry (BLI). Each mAb was immobilized on a protein-A sensor and incubated with each of the rRBD mutants [N501Y, S477N, E484K, N439K, K417N and Y453F] or WT rRBD as a control, for 180 sec.

### Comparison of binding to the SARS-CoV-2 spike protein by MD65 and the commercial licensed LY-CoV555 mAbs

Amongst the four RBD-specific mAbs studied here, MD65 is the most effective antibody in terms of *in vitro* neutralization. Additionally, *in vivo* studies demonstrated that MD65 effectively elicited post-exposure protection in mice at relatively low doses (Rosenfeld et al., 2021). MD65 (whose variable regions are encoded by the IGHV3-66 and IGKV3-20 germline heavy and light chain alleles, respectively), belongs to a public clonotype (frequently encoded by IGHV3-53/3-66) that was extensively characterized in the context of SARS-CoV-2 neutralizing human antibodies (Barnes et al., 2020; Fagiani et al., 2020; Tan et al., 2021; Yuan et al., 2020), specifically targets the receptor binding motif, competing with hACE2 binding. Noteworthy, recent studies showed that binding and neutralization by antibodies belonging to this public clonotype are weakened by either the K417N or E484K replacements (Andreano et al., 2020; Yuan et al., 2021). Specifically, the E484K mutation, which is present in several SARS-CoV-2 natural isolates (including the B.1.351, P.1 and recently identified “B.1.1.7+E484K” VOCs), was reported to be associated with lower susceptibility to neutralization by some mAbs, convalescent plasma and sera collected from vaccinated individuals (Chen et al., 2021b; Wang et al., 2021a; Wang et al., 2021b).

The RBD-specific therapeutic mAb LY-CoV555 [Bamlanivimab; (Chen et al., 2021a)], encoded by the germline alleles: IGHV1-69; IGKV1-39, was also shown to block hACE2 binding by SARS-CoV-2 (Jones et al., 2021), and accordingly, possibly compete with MD65. However, although they may target close epitopes, their recognition pattern may differ. In order to test whether LY-CoV555 functionality is affected in a similar manner as MD65, the commercially available LY-CoV555 mAb was used in binding experiments. First, the binding profile of the LY-CoV555 was tested by ELISA against the SARS-CoV-2 spike protein, and compared to that of MD65 (Figure 3A), demonstrating similar binding profiles. An epitope-binning experiment using BLI was performed in the presence of MD65 and MD29 (as a negative control). As shown in Figure 3B, LY-CoV555 failed to bind the rRBD protein, presented in complex with MD65 mAb, while significant binding was observed when the rRBD was presented in complex with MD29 (which binds a different epitope than MD65). These results clearly indicate that LY-CoV555 and MD65 target overlapping epitopes. Next, the LY-CoV555 binding capacity towards the panel of RBD mutants was evaluated (Figure 3C), demonstrating equivalent binding of rRBD-N439K, Y453Y, S477N and N501Y, compared to the WT rRBD. However, LY-CoV555 binding was completely obstructed by the E484K substitution. This observation is in line with recently reported studies suggesting that the E484K substitution is accountable for the abolishment of neutralization of SARS-CoV-2 natural variants that carry this mutation by LY-CoV555 mAb (Starr et al., 2021b; Wang et al., 2021a).

**Figure 3.**
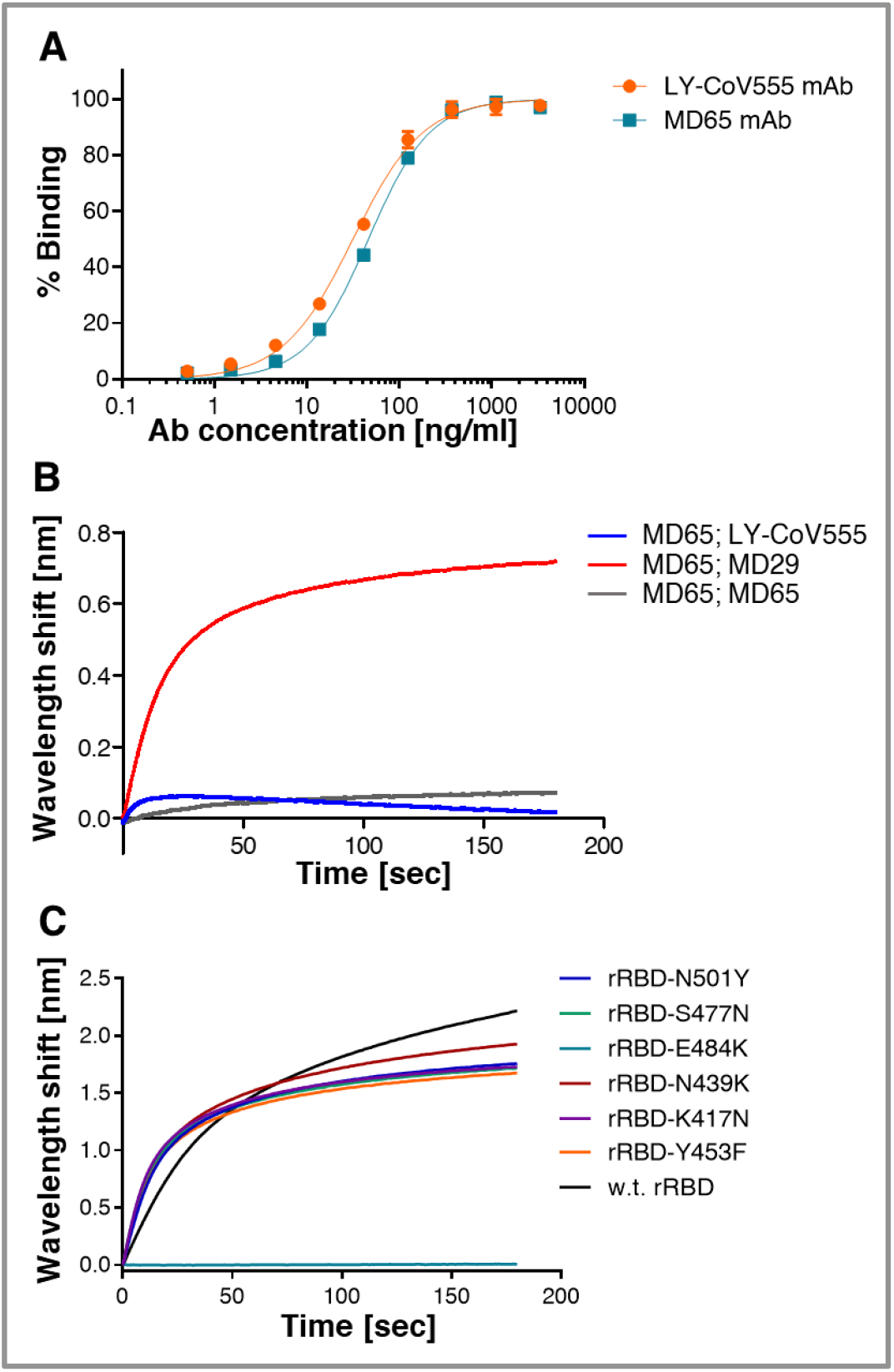
Binding characterization of the LY-CoV555 mAb (Bamlanivimab). **A.** Binding of the LY-CoV555 mAb was evaluated by ELISA against SARS-CoV-2 spike protein. MD65 was included for comparison. Data shown represent average of triplicates ±SEM **B.** BLI was applied for epitope binning experiments. MD65 antibody was biotinylated, immobilized on a streptavidin sensor and saturated with WT rRBD protein. The complex was then incubated for 180 s with LY-CoV555 or MD29 and MD65 as controls. Time 0 represents the binding to the MD65-rRBD complex. **C.** LY-CoV555 binding of the indicated single-mutated rRBDs and the WT rRBD was evaluated by BLI. LY-CoV555 antibody was immobilized on a protein-A sensor and incubated with each indicated rRBD for 180 sec.

### Structural modeling of anti-RBD antibodies

To define the molecular basis for the observed high cross-reactivity of MD65 mAb against all inspected mutant RBDs, we modeled its variable domain structure using *AbPredict2* [(Lapidoth et al., 2019); Figure 4]. *AbPredict2* uses Rosetta energy calculations as the sole criterion to predict a model structure on the basis of the variable domain sequence, ignoring sequence homology to existing antibody structures (Norn et al., 2017), producing energy-relaxed models. We noted that the recently solved structure of antibody CovA2-04 in complex with the RBD [Protein Data Bank entry 7jmo; (Wu et al., 2020a)] is very similar to the top-ranked *AbPredict2* model of MD65 (Cα and carbonyl oxygen root mean square deviation < 1.0 Å). The two antibodies are also highly similar in their primary amino-acid sequence (93% V gene sequence identity; Figure 4A), and both derive from the same heavy and light-chain germline genes (IGHV3-53/66 and IGKV3-20, respectively), hence are assigned to the same public clonotype. The differences between the two antibodies are mostly restricted to a diverged amino acid residue in L1, a deletion in L3, and different H3 sequences. The high sequence and structure similarity suggests that the bound state observed for CovA2-04 may provide a reliable structural framework for analyzing MD65 binding to RBD variants. Therefore, the MD65 model structure was aligned to the CovA2-04 structure (Figure 4B) to obtain a model of the interaction of MD65 with RBD.

**Figure 4:**
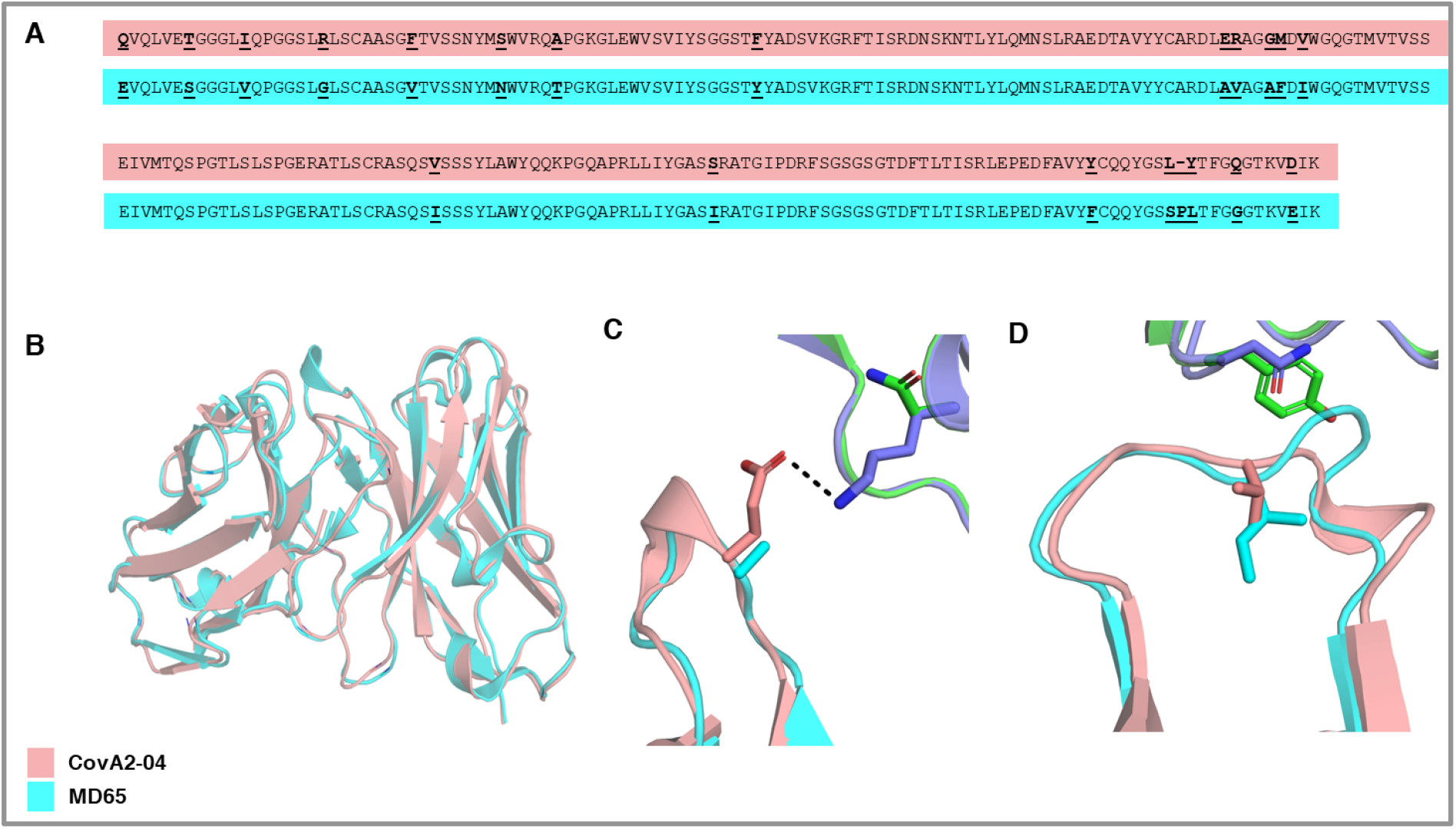
Structural basis of MD65 binding to COVID-19 spike variants determined by comparative modeling to the CovA2-04 mAb. **A.** Alignment of the primary amino-acid sequences of the heavy (2 upper sequences) and light (2 lower sequences) chain variable domains of the two antibodies compared (CovA2-04 in pink and MD65 in cyan). Diverged residues are indicated by bold and underlined letters. **B-D.** MD65 model structure (cyan) aligned with CovA2-04 crystal structure (pink); WT (PDB entry 7jmo) and B.1.351 spike (PDB entry 7nxa); violet and green, respectively). **B.** A view of the superimposed variable domain models of the two mAbs, indicating the close correspondence of the MD65 model structure and the experimentally determined structure of CovA2-04. **C.** The Lys residue at position 417 of the WT spike protein forms a stabilizing hydrogen bond (dashed line) with the Glu residue at position 100 of CovA2-04. The K417N present in B.1.351 abrogates this stabilizing interaction leading to potential strain in binding to the negative surface on the CDR H3 loop of CovA2-04. The Ala at the analogous position in MD65 may relieve this strain. **D.** The small-to-large N501Y substitution on the B.1.351 spike may physically overlap with the CDR L1 of CovA2-04. The V29I difference in MD65 (compared to CovA2-04) modifies the CDR L1 backbone conformation, expanding the space in this region for the bulkier Tyr residue of the spike.

We initially focused on the K417N mutation located on the B.1.351 spike protein (Figure 4C). Despite the high sequence homology of CovA2-04 and MD65, this mutation abrogates binding to the former while minimally perturbing binding to the latter (Yuan et al., 2021). Whereas CDR H3 in CovA2-04 presents a negatively charged sidechain (Glu100; annotated as Glu97 according to Kabat numbering scheme) to counter the positive charge on the Lys at position 417 in RBD, the model shows that the H3 loop of MD65 is neutral. Thus, the K417N mutation may lead to electrostatic strain in binding the CovA2-04 antibody but not in that of MD65. Negative charges are also observed in this position in other RBD-binding antibodies derived from the IGHV3-53/66 germline gene (Yuan et al., 2021) and these may be similarly impacted by the K417N mutation.

Structural modeling may also provide an explanation for the ability of MD65 to neutralize RBD variants that exhibit the N501Y mutation (Figure 4D). RBD position 501 is proximal to the tip of CDR L1 of CovA2-04. The *AbPredict2* model predicts that the CDR L1 adopts a different backbone conformation than CovA2-04 due to the CDR L1 mutation V29I (refers to position 28 according to the Kabat numbering scheme). In this altered backbone conformation, the CDR L1 of MD65 provides more space for the bulky Tyr at RBD position 501. Finally, the structure model suggests that RBD Glu at position 484 is distant from the interaction with MD65. Therefore, even radical mutations at this position would not impact antibody binding, as is indeed observed in the binding and neutralization experiments. Thus, it may be concluded that electrostatic strain at CDR H3 and steric hindrance in CDR L1 provide a likely mechanistic basis for understanding the differential effects of RBD mutations on binding affinity and neutralization in this class of antibodies. It remains to be seen whether this explanation extends to additional antibodies belonging to this class and other spike variants that emerge in the future.

### Binding SARS-CoV-2 multiply mutated-S1 versions by specific mAbs

The retained binding capabilities, observed for the tested mAbs towards the individual RBD mutations, may not necessarily predict their interaction in the context of multiple mutations, present in emerged VOCs. Therefore, we studied the ability of the four RBD-specific mAbs to bind recombinant mutated spike S1 subunit proteins, representing the RBD accumulated mutations associated with the B.1.1.7 and B.1.351 genetic variants (schematically depicted in Figures 1B and 1C, respectively). BLI analyses (Figure 5) were applied for binding evaluation of recombinant S1:Δ 69-70; Δ 144; N501Y; A570D; D614G; P681H, representing all the modifications encountered in the S1 of the B.1.1.7 genetic variant (Figure 5, B.1.1.7 rS1, blue curves) and of recombinant S1:K417N; E484K; N501Y; D614G, representing the RBD-related substitutions of B.1.351 (B.1.351 rS1; red curves). Recombinant WT S1 subunit (WT rS1; black curves) was used for comparison. While binding of B.1.1.7 rS1 by MD65, MD29, BL6 and LY-CoV555 was not impaired by the mutations (Figures 5A, 5C, 5D and 5E, respectively), MD62 binding was reduced by ~50% (Figure 5B). This observation may not be attributed to the minor reduction (8% loss of binding) observed in binding of N501Y-RBD mutant by this antibody (Figure 2B) as it represents the only substitution in the RBD of B.1.1.7. Therefore, it can be speculated that structural changes in the B.1.1.7 rS1 (Figure 1B) involving the mutated NTD, allosterically affected MD62 binding.

**Figure 5.**
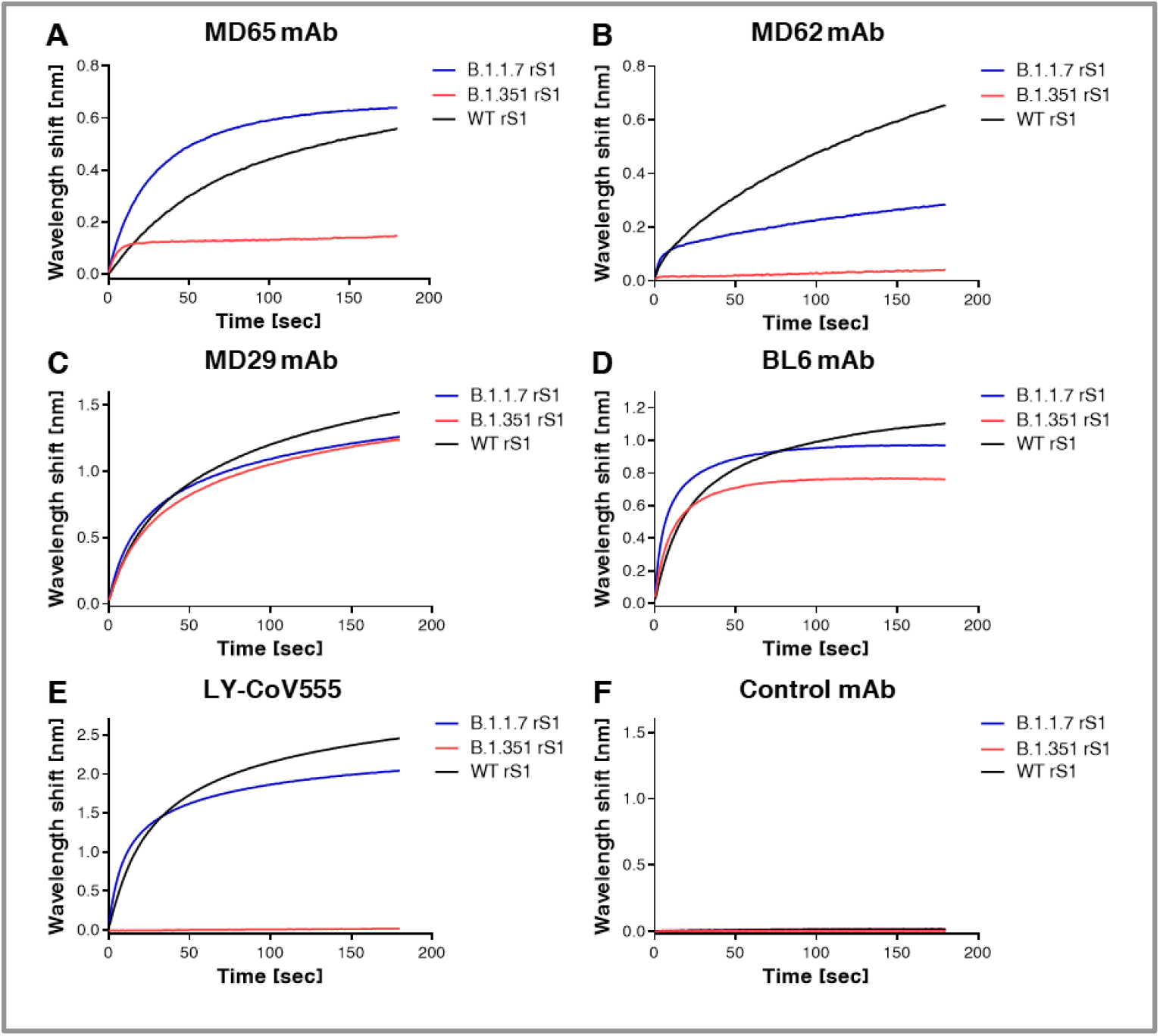
Binding of rS1 variants by RBD-specific mAbs. BLI was applied to evaluate the ability of each tested mAb to bind the indicated recombinant multiply-mutated spike S1 subunit proteins: B.1.1.7 rS1 [Δ 69-70; Δ 144; N501Y; A570D; D614G; P681H] and B.1.351 rS1 [K417N; E484K; N501Y; D614G]. Each of the indicated mAbs (**A-E**), was immobilized on protein-A sensor and incubated for 180 sec with B.1.1.7 (blue) or B.1.351 (red) rS1 variants or with the WT rS1 (black). Non-specific control antibody (anti-ricin MH75) was included (**F**). The figure includes representative graphs of at least two independent repeats of each experiment, yielding similar results.

The B.1.351 rS1 (schematically depicted in Figure 1C) includes the N501Y substitution, as well as the K417N and E484K replacements that were previously shown to substantially impair binding by various mAbs (Chen et al., 2021b; Cheng et al., 2021; Starr et al., 2021b; Wang et al., 2021b). Thus, in line with the individual mutation binding results (see Figure 3C for LY-CoV555 and Figure 2D for BL6), the observed binding abrogation of the B.1.351 rS1 by LY-CoV555 mAb (Figure 5B) and the mild (18%) reduction observed for BL6 (Figure 5D) can be attributed mainly to the E484K substitution. Similarly, the complete loss of binding by MD62 (Figure 5B) and significant loss of binding by MD65 (~65%; Figure 5A), may have been mediated by the K417N substitution (see Figures 2B and 2A for MD62 and MD65, respectively).

### Antibody-mediated neutralization evaluated by a cell-culture plaque reduction test

In order to conclusively determine the potential of the B.1.1.7 and B.1.351 SARS-CoV-2 genetic variants (see Supplementary Figure 1, for their spike mutations) to escape immune-neutralization, we evaluated the ability to countermeasure the B.1.1.7 and B.1.351 live variants of the four RBD-specific mAbs, LY-CoV555, and two additional mAbs, targeting separate epitopes of the NTD (BLN14 and BLN12; (Noy-Porat et al., 2021); see Supplementary Figure 3 for data pertaining to these 2 anti-NTD antibodies to B.1.1.7 rS1). To this end, a plaque reduction neutralization test (PRNT) was applied in which each mAb was tested against either the B.1.1.7 or B.1.351 variants or the parental WT SARS-CoV-2 strain.

The results, presented in Figure 6, indicated an effective neutralization of the B.1.1.7 variant (blue curves) by anti-RBD MD65 (panel A), MD62 (panel B), MD29 (Panel C), BL6 (Panel D) as well as LY-CoV555 (panel E), a neutralization that was not impaired compared to the WT SARS-CoV-2 strain (black curves). The calculated IC_50_ values, characterizing the neutralization potency of each inspected antibody with respect to the three tested viral strains, are tabulated in Figure 6H. Interestingly, although MD62 revealed reduced binding to the B.1.1.7 rS1 compared to the WT rS1 (Figure 5B), its neutralization capacity of both viral strains was commensurate (Figure 6B). Overall, it can be concluded that all anti-RBD mAbs studied here fully retained their potency towards the B.1.1.7 variant.

**Figure 6.**
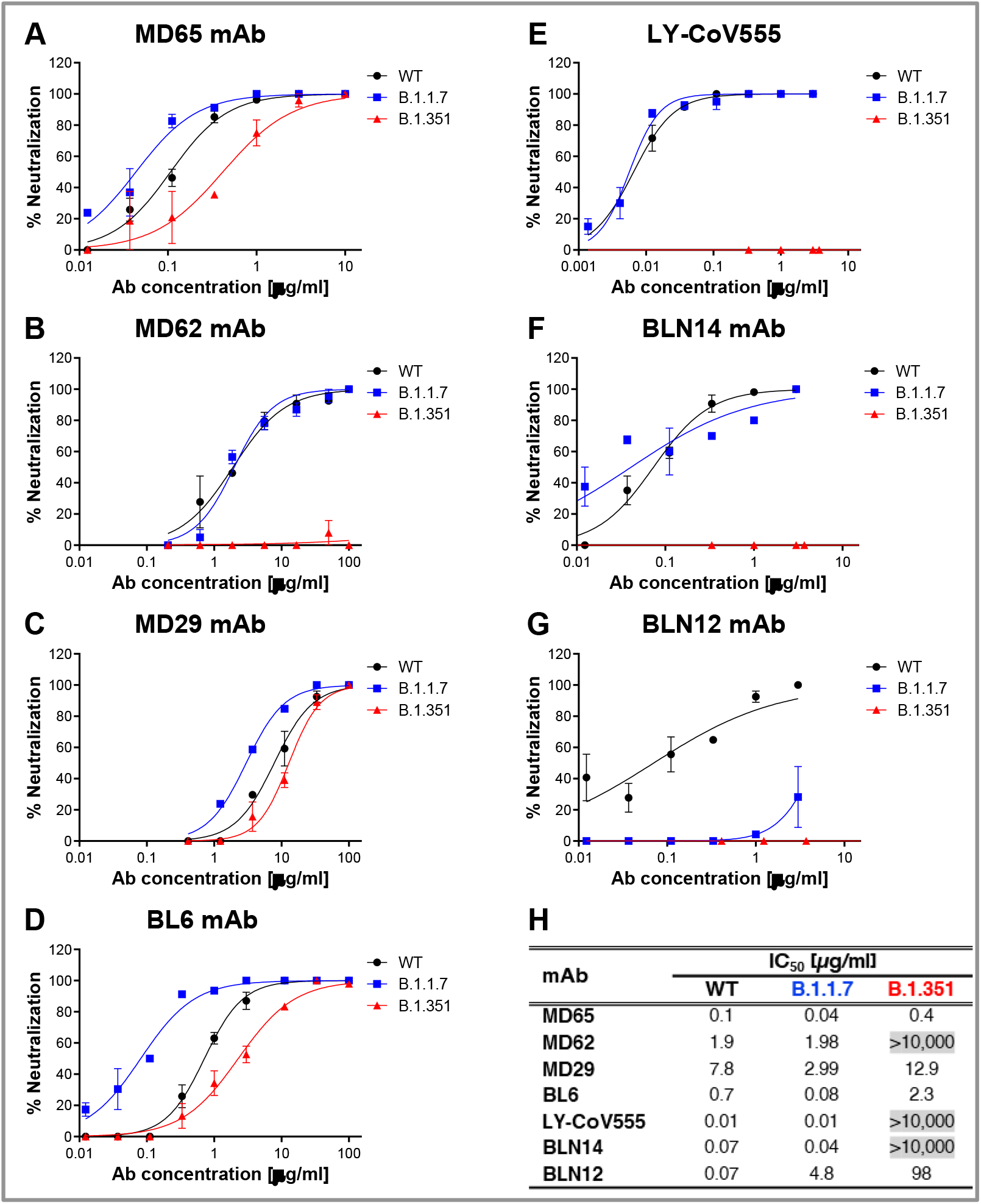
Neutralization of SARS-CoV-2 B.1.1.7 and B.1.351 by RBD and NTD-specific mAbs. Neutralization capacity of the RBD-specific mAbs: MD65 (**A**), MD62 (**B**), MD29 (**C**), BL6 (**D**) and LY-CoV555 (**E**), and of the NTD-specific BLN14 (**F**) and BLN12 (**G**), was evaluated by plaque reduction neutralization test (PRNT). The *in vitro* neutralization of each of the listed mAbs was assessed against both SARS-CoV-2 B.1.1.7 (blue) and B.1.351 (red) variant, compared to WT SARS-CoV-2 strain (black). Neutralization potency was determined by the ability of each antibody (at indicated concentrations) to reduce plaque formation. Results are expressed as percent inhibition of control without Ab. The figure includes a representative graphs of at least two independent repeats of each experiment, yielding similar results. **H**. Summary of the calculated IC_50_ values [μg/ml]. IC_50_ >10,000 indicates complete loss of neutralization capacity, emphasized by gray shading. The neutralization results, together with previously published biochemical data of the six inspected mAbs are summarized in Supplementary Table 1.

As could be anticipated, the B.1.351 variant (Figure 6, red curves) manifested a higher immune escape potential compared to the B.1.1.7 variant. In line with the complete loss of binding of the respective viral rS1 by MD62 (Figure 5B) and LY-CoV555 (Figure 5E), the neutralization capacity of the two mAbs (MD62 and LY-CoV555) against the B.1.351 variant, was completely abolished (Figures 6B and 6E, respectively). By contrast, MD65, MD29 and BL6 effectively neutralized the B.1.351 variant (Figures 6A, 6C and 5D), albeit, with a partial decrease in potency.

Two anti-NTD mAbs, previously shown to potently neutralize SARS-CoV-2, were also tested in the *in vitro* neutralization assay. The two mAbs, BLN14 and BLN12, differed in their potency against the B.1.1.7 variant (Figures 6F and 6G, respectively), with BLN14 showing comparable neutralization to that of the WT, while BLN12 neutralization activity was markedly hampered. These results are consistent with binding experiments showing strong binding of B.1.1.7 rS1 by BLN14 and no binding by BLN12 (Supplementary Figure 3). Epitope mapping previously revealed that BLN12 binds a linear epitope which resides between amino acids 141-155 and also recognizes an N-glycan at position 149 (Noy-Porat et al., 2021). It can therefore be speculated that the deletion of a Tyr residue at position 144 in the B.1.1.7 variant is responsible for the loss of neutralization of this mAb. BLN14 recognizes a conformational epitope which apparently was not significantly altered by the Y144 deletion. However, the neutralization capability of both mAbs was dramatically reduced in the case of the B.1.351 variant, suggesting a considerable structural change at its NTD. This observation is in agreement with previous studies, which indicated a frequent loss of functionality among NTD-specific mAbs (Andreano et al., 2020; Wang et al., 2021b), especially towards variants containing modifications in NTD supersite (Cerutti et al., 2021; McCallum et al., 2021) associated with significant structural alterations.

### Antibody-mediated protective value evaluated by post-exposure administration in a transgenic murine model of COVID-19

In line with the *in vitro* neutralization performance of the studied mAbs, MD65, BL6 and BLN14 mAbs were selected for further assessment of their therapeutic potential *in vivo* against various SARS-CoV-2 variants. Accordingly, K18-hACE2 transgenic mice were intranasally infected with a lethal dose of either one of the SARS-CoV-2 tested strains. The infection was characterized by body weight loss accompanied with high mortality (70-100%, i.e. animals administered with PBS; Figure 7, gray lines) within 6-12 days following infection. One mg of each of the mAb, was administered intraperitoneally (IP) two days post infection (dpi). Body weight and survival of experimental animals were monitored daily for 21 days. The data depicted in Figure 7 clearly demonstrate that antibody administration to infected animals results in a strong protective effect against the three variants [see also (Noy-Porat et al., 2021; Rosenfeld et al., 2021) for the effect of the MD65 and BLN14 antibodies against the WT version of SARS-CoV-2]. Notably, the RBD-specific antibodies MD65 and BL6 exhibited substantial therapeutic ability against all three viral variants inspected. Treatment with BL6 rescued 83-100% of infected animals regardless of the infective strain. MD65 afforded complete protection against the WT and B.1.1.7 strains and 50% protection against the B.1.351 strain. This apparently lower extent of protection is substantial in view of the fact that a considerably higher dose of this variant was administrated to the experimental animals (see Method Details section). The anti-NTD antibody BLN14 demonstrated high potency against the WT and B.1.1.7 variant yet, post-exposure treatment with BLN14 mAb failed to rescue mice from infection with the B.1.351 strain. Taken together, these results are in good agreement with the *in vitro* BLI binding and PRNT data establishing that these *in vitro* tests may provide a reliable predictive means for evaluation of therapeutic antibodies.

**Figure 7.**
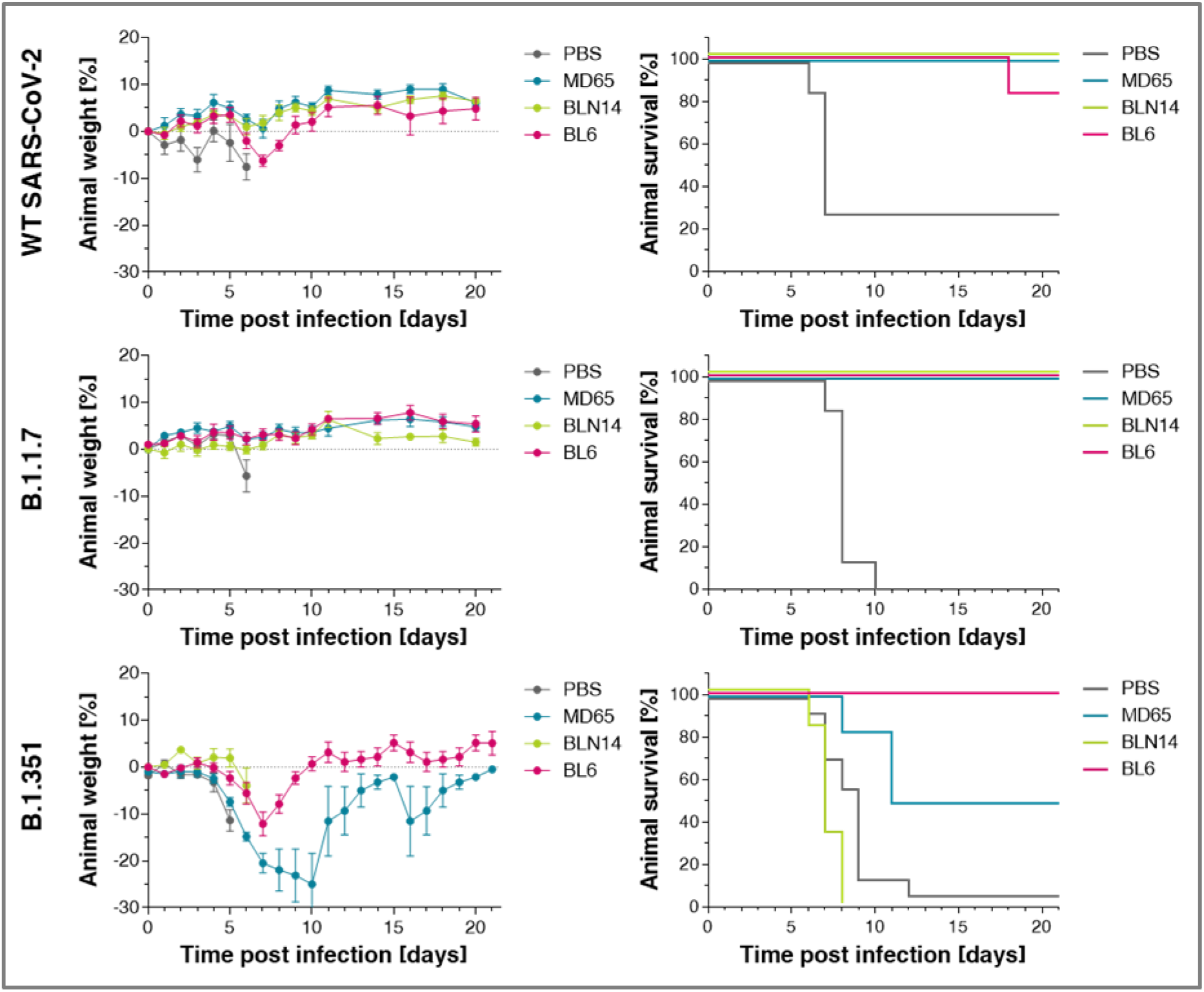
Post-exposure therapeutic potency of MD65, BL6 and BLN14 in K18-hACE2 mice infected with various SARS-CoV-2 strains. The SARS-CoV-2 strains are indicated in the left vertical lane and the administered antibodies (colored differently) in the legends within each individual panel. A single dose of 1 mg Ab/animal of each indicated mAb was administered at day 2 post viral infection. Left panels: body weight profiles. Body weight is displayed as percentage change of initial weight. Only data of the first six days is presented in the control group exhibiting significant mortality. Right panels: Kaplan-Meyer surviving curves.

## Conclusions

The unprecedented scale of the COVID-19 pandemic, combined with selective pressure for escaping immune responses, boosted the rapid evolution of SARS-CoV-2 virus resulting in antigenic variability which might jeopardize the potency of pre- and post-exposure immunotherapies. Consequently, attention must be given to the development of mAb treatments that may combat emerging variants. In this perspective, it is of high importance to re-evaluate anti-SARS-CoV-2 mAbs previously shown to exhibit therapeutic potential against the original version of the virus. Furthermore, the impact of individual mutations on the neutralization potency of mAbs may provide important information impacting the preparedness for future anticipated antigenic drifts of the virus. In the current report, we document the neutralization of the most abundant B.1.1.7 variant (as of today) by four anti-RBD and one anti-NTD mAbs that we recently generated and determined their therapeutic potential against the original version of the virus. Furthermore, three RBD-specific mAbs (MD65, MD29 and BL6), retained neutralization against the B.1.351 VOC. These findings are supported by binding experiments conducted with individual and combined mutations derived from various variants. The binding and neutralizing data of the six inspected mAbs targeting distinct epitopes within S1 are summarized in Supplementary Table 1. The E484K and K417N substitutions in the RBD reported to mediate the lower susceptibility to neutralization by a significant proportion of reported mAbs, including clinically-used LY-CoV555 and REGN10933 as well as by immune post-vaccination sera (Chen et al., 2021b; Cheng et al., 2021; Dejnirattisai et al., 2021; Hu et al., 2021; Rees-Spear et al., 2021; Starr et al., 2021b; Wang et al., 2021a; Wang et al., 2021b; Yuan et al., 2021). Of note, the anti-RBD antibody MD65, shown here to retain its neutralizing potential against emerging variants, was recently suggested by extensive pre-clinical studies to be an important therapeutic for efficient clinical intervention in COVID-19 cases (Rosenfeld et al., 2021). Structural modeling identified specific residues in the sequence of the MD65 mAb which may explain the potency of this antibody against all the viral variants inspected. Finally, the binding and neutralizing data were confirmed by *in vivo* protection of infected transgenic mice administered with the various antibodies two days post-infection.

In conclusion, the present study substantiates the ability of recently reported mAbs to serve, individually or in combination as a cocktail formulation, for designing therapeutic approaches efficient against emerging SARS-CoV-2 variants.

## Acknowledgments

We thank Prof. Dr. Christian Drosten at the Charité Universitätsmedizin, Institute of Virology, Berlin, Germany for providing the SARS-CoV-2 BavPat1/2020 strain. We wish to express our gratitude to our colleagues Dr. Sharon Melamed, Boaz Politi, Dr. Liat Bar-On, Yfat Yahalom-Ronen and Dr. Emanuelle Mamroud, for fruitful discussions and support. The graphical abstract was created with BioRender.com. Research in the Fleishman lab was supported by the Weizmann Institute CoronaVirus Fund and by a charitable donation in memory of Sam Switzer.

## Author Contributions

E.M., A.Z., R.A., T.N-P., E.P., A.M., Y.L., E.E., A.T., M.A., D.G., S.J.F., O.M. and R.R. designed, carried out and analyzed the data. M.M., E.M, N.Z., I.N., and L.K isolated, sequenced and provided the SARS-CoV-2 variant strains. N.P., H.T. and T.I. cultured and prepared SARS-CoV-2 viruses for the neutralization experiments. O.Z. and S.W. provided crucial reagents. S.Y., S.C.S. added fruitful discussions. R.R., O.M, T.C., A.Z., and T. N-P wrote the manuscript. O.M. and R.R. supervised the project. All authors have reviewed and approved the final manuscript.

## Competing Interests

Patent application for the described antibodies was filed by the Israel Institute for Biological Research. None of the authors declared any additional competing interests.

## STAR Methods

### RESOURCE AVAILABILITY

#### Lead contact

Further information and requests for resources and reagents should be directed to and will be fulfilled by the Lead Contact, Ohad Mazor from the Israel Institute for Biological Research.

#### Materials availability

Antibodies are available for research purposes only under an MTA, which allows the use of the antibodies for non-commercial purposes but not their disclosure to third parties. All other data are available from the Lead contact upon reasonable requests.

#### Data and code availability

The published article includes all dataset generated or analyzed during this study.

### EXPERIMENTAL MODELS AND SUBJECT DETAILS

#### Animals

Female K18-hACE2 transgenic (B6.Cg-Tg (K18-hACE2)2Prlmn/J HEMI; Jackson Laboratories, USA) mice, age 8-16 weeks, were maintained at 20-22°C and a relative humidity of 50 ± 10% on a 12 hours light/dark cycle, fed with commercial rodent chow (Koffolk Inc.) and provided with tap water ad libitum. Treatment of animals was in accordance with regulations outlined in the U.S. Department of Agriculture (USDA) Animal Welfare Act and the conditions specified in the Guide for Care and Use of Laboratory Animals, National Institute of Health, 2011. Animal studies were approved by the local ethical committee on animal experiments (protocol number M-57-20).

#### Cells and virus strains

ExpiCHO-S (Thermoscientific, USA, Cat# A29127) were used for expression of recombinant proteins as described above.

Vero E6 (ATCC^®^ CRL-1586™) were obtained from the American Type Culture Collection. Cells were grown in Dulbecco’s modified Eagle’s medium (DMEM) supplemented with 10% fetal bovine serum (FBS), MEM non-essential amino acids (NEAA), 2 mM L-glutamine, 100 Units/ml penicillin, 0.1 mg/ml streptomycin and 12.5 Units/ml Nystatin (P/S/N) (Biological Industries, Israel). Cells were cultured at 37°C, 5% CO_2_ at 95% air atmosphere.

Wild type (WT) SARS-CoV-2 strain (GISAID accession EPI_ISL_406862) was kindly provided by Bundeswehr Institute of Microbiology, Munich, Germany.

WT SARS-CoV-2, isolate Human 2019-nCoV ex China strain BavPat1/2020, was kindly provided by Prof. Dr. Christian Drosten (Charité, Berlin, Germany) through the European Virus Archive – Global (EVAg Ref-SKU:026V-03883).

SARS-CoV-2 B.1.1.7 (501Y.V1) variant was isolated on Dec 2020 from a person who came back from the UK. The identity of the B.1.1.7 strain was confirmed using NGS. SARS-CoV-2 B.1.351 (501Y.V2) variant was isolated on Jan 2021 from a person who was in contact with a patient who came back from South Africa. The identity of the B.1.351 strain was confirmed using NGS.

Stocks were prepared by infection of Vero E6 cells for two days. When viral cytopathic effect (CPE) was observed, media were collected, clarified by centrifugation, aliquoted and stored at −80°C. Titer of stock was determined by plaque assay using Vero E6 cells.

Handling and working with SARS-CoV-2 was conducted in BL3 facility in accordance with the biosafety guidelines of the IIBR.

### METHOD DETAILS

#### Recombinant Proteins

The SARS-CoV-2 spike (S) stabilized soluble ectodomain, S1 subunit (WT rS1) and receptor binding domain (WT rRBD) were produced as previously described (Noy-Porat et al., 2020).

The following His-tagged recombinant proteins were purchased from Sino Biologicals: B.1.1.7 rS1-SARS-CoV-2 spike S1 [Δ 69-70; Δ 144; N501Y; A570D; D614G; P681H], cat#40591-V08H12; B.1.351 rS1-SARS-CoV-2 spike S1 [K417N; E484K; N501Y; D614G], cat#40591-V08H10; spike RBD[N501Y] cat#40592-V08H82; spike RBD[S477N] cat#40592-V08H46; spike RBD[E484K] cat#40592-V08H84; spike RBD[N439K] cat#40592-V08H14; spike RBD[K417N] cat#40592-V08H59; spike RBD[Y453F] cat#40592-V08H80.

All antibodies (except LY-CoV555) were produced as full IgG1 antibodies as described (Barlev-Gross et al., 2021; Rosenfeld et al., 2021), expressed using ExpiCHO™ Expression system (Thermoscientific, USA) and purified on HiTrap Protein-A column (GE healthcare, UK). The integrity and purity of the antibodies were analyzed using SDS-PAGE. Isolation and characterization of the MD29, MD65 and MD62 mAbs, targeting epitopes I-III on the RBD as previously reported. The BL6 mAb was isolated as described (Barlev-Gross et al., 2021) and is representing epitope IV on the RBD (competing with the MD47 mAb (Noy-Porat et al., 2020). BLN12 and BLN14 mAbs, targeting two distinct epitopes on the NTD, as previously reported (Noy-Porat et al., 2021).

LY-CoV555 (Bamlanivimab) (~2.5 mg Ab/ml in 0.9% Sodium Cholride), was obtained as a remnant from an infusion bag and its set following administration to a COVID-19 patient at Kaplan Medical Center.

All antibodies were extensively dialyzed against PBS and filter-sterilized prior to any *in vitro* or *in vivo* experimentation.

### ELISA

Direct ELISA (Noy-Porat et al., 2016) consisted of coating microtiter plates with 1 μg/ml of recombinant SARS-CoV-2 spike. ELISA was applied with AP-conjugated Donkey anti-human IgG (Jackson ImmunoResearch, USA, Cat# 709-055-149 lot 130049; used at 1:2000 working dilution) following detection using *p*-nitrophenyl phosphate (*p*NPP) substrate (Sigma, Israel).

#### Biolayer interferometry (BLI)

Binding studies were carried out using the Octet system (ForteBio, USA, Version 8.1, 2015) that measures biolayer interferometry (BLI). All steps were performed at 30°C with shaking at 1500 rpm in a black 96-well plate containing 200 μl solution in each well. For assessment of binding to S1 variants or mutated RBD, antibodies were captured on Protein-A or anti-Fab CH1 sensors (FAB2G) and incubated with recombinant S1 (WT, B.1.1.7 or B.1.351) or recombinant RBD (WT or mutated) at a constant concentration of 10 μg/ml for 180 sec and then transferred to buffer containing wells for additional 60 sec. Binding was measured as changes over time in light interference. Parallel measurements from unloaded biosensors were used as control. The anti-ricin MH75 mAb, used as isotype control (Figure 5F). For the comparison of the binding capacity of each tested mAb to a constant concentration of recombinant RBD or S1, the area under curve (AUC) was calculated for each binding curve, using GraphPad Prism 5, and percent binding was calculated compared to the WT protein, representing 100% binding.

For epitope binning, MD65 antibody was biotinylated, immobilized on streptavidin sensor, incubated with a fixed concentration of WT rS1 (20 μg/ml) to reach saturation, washed and incubated with non-labeled LY-CoV555 for 180 sec. MD29 and MD65 were used as positive and negative controls, respectively.

#### Plaque reduction neutralization test (PRNT)

Plaque reduction neutralization test (PRNT), performed essentially as described (Yahalom-Ronen et al., 2020).Vero E6 cells were seeded overnight at a density of 0.5e6 cells/well in 12-well plates. Antibody samples were 3-fold serially diluted (ranging from 200 to 0.002 μg/ml) in 400 μl of MEM supplemented with 2% FBS, MEM non-essential amino acids, 2 mM L-glutamine, 100 Units/ml penicilin, 0.1 mg/ml streptomycin and 12.5 Units/ml Nystatin (Biological Industries, Israel). 400 μl containing 300 PFU/ml of each SARS-CoV-2 strain, were then added to the mAb solution supplemented with 0.25% guinea pig complement sera (Sigma, Israel) and the mixture incubated at 37°C, 5% CO_2_ for 1 h. Two hundred μl of each mAb-virus mixture was added in duplicates to the cells for 1 h. Virus mixture w/o mAb was used as control. 2 ml overlay [supplemented MEM containing 0.4% tragacanth (Sigma, Israel)] were added to each well and plates were further incubated at 37°C, 5% CO_2_ for 48 h for WT and B.1.351 strains or 5 days for the B.1.1.7 strain. The number of plaques in each well was determined following media aspiration, cells fixation and staining with 1 ml of crystal violet (Biological Industries, Israel). Half-maximum inhibitory concentration (IC_50_) was defined as mAb concentration at which the plaque number was reduced by 50%, compared to plaque number of the control (in the absence of Ab).

### Animal experiments

All animal experiments involving SARS-CoV-2 were conducted in a BSL3 facility. Infection experiments were carried out using SARS-CoV-2 BavPat1/2020 (WT), B.1.1.7 and B.1.351 strains. SARS-CoV-2 virus diluted in PBS supplemented with 2% FBS (Biological Industries, Israel) was used to infect anesthetized mice by intranasal instillation. For mAbs protection evaluation, mice were treated intraperitoneally by single administration of 1 ml volume, containing 1 mg Ab/mouse, two days following infection with 500, 10 or 10,000 PFU of the WT, B.1.1.7 and B.1.351 SARS-CoV-2 strains, respectively. Control groups were administered with PBS. Body weight was monitored daily throughout the follow-up period post infection.

#### Antibody structure prediction

Antibody structure prediction was done using *AbPredict2* with default settings. RMSD calculation and alignments were done using Pymol. *AbPredict2* is available for academic use at http://AbPredict.weizmann.ac.il.

### QUANTIFICATION AND STATISTICAL ANALYSIS

All Biolayer Interferometry assays were analyzed using the Octet Data analysis software (ForteBio, Version 8.1) and visualized using GraphPad Prism 5.

ELISA results were analyzed using GraphPad Prism 5. Mean and SEM were calculated where appropriate and are presented in the relevant figures.

All the following statistical analyses were conducted using GraphPad Prism 5. For *In vitro* neutralization experiments, mean and SEM were calculated for each concentration, curves were fitted using nonlinear regression and IC50 values were extrapolated from the resulting curves. Results are presented in the relevant figures, in the results section and in Supplementary Table 1.

Area under curve (AUC) was calculated for each binding curve presented in Figure 5. Results are presented in Supplementary Table 1.

For all experiments, exact value and meaning of n are presented in the figure legends.

For antibody structure modeling, the amino acid sequence of the variable domain of MD65 was submitted to the *AbPredict2* webserver which is available freely for non-commercial use (http://AbPredict.weizmann.ac.il).

## Notes

### Summary of Updates

Revision of the text Change in the authors list

